# Meta-16S rRNA gene phylogenetic reconstruction reveals astonishing diversity of the cosmopolitan myxobacteria

**DOI:** 10.1101/754119

**Authors:** Yang Liu, Qing Yao, Hong-Hui Zhu

**Affiliations:** State Key Laboratory of Applied Microbiology Southern China, Guangdong Provincial Key Laboratory of Microbial Culture Collection and Application, Guangdong Open Laboratory of Applied Microbiology, Guangdong Microbial Culture Collection Center (GDMCC), Guangdong Institute of Microbiology, Guangdong Academy of Sciences, Guangzhou 510070, PR China; College of Horticulture, South China Agricultural University, Guangdong Province Key Laboratory of Microbial Signals and Disease Control, Guangdong Engineering Research Center for Grass Science, Guangdong Engineering Center for Litchi, Guangzhou 510642, China

**Keywords:** myxobacteria, phylogenetic diversity, geographical distribution, environments

## Abstract

Numerous ecological studies for myxobacteria have been conducted well, but their true diversity hidden in plain sight remains to be explored. To bridge this gap, we here implemented a comprehensive survey of diversity and distribution of myxobacteria by using 4997 publicly available 16S rRNA gene sequences (≥1200 bp) collected from several hundreds of sites across multiple countries and regions. In the study, the meta-16S rRNA gene phylogenetic reconstruction clearly revealed that these sequences were classified to 998 species, 445 genera, 58 families, and 20 suborders, highlighting a considerable taxonomic diversity of myxobacteria, the great majority of which belonged to new taxa. Most cultured myxobacteria including the well-described type strains were strongly inclined to locate on the shallow branches of the phylogenetic tree; on the contrary, the majority of uncultured myxobacteria the deep branches. The geographical analysis of sequences based on their environmental categories clearly demonstrated that myxobacteria showed a nearly cosmopolitan distribution, despite the presence of some habitat-specific taxa, especially at genus and species levels. Among all abundant suborders, members of Suborder_4, Suborder_15, and Suborder_17 were more widely distributed in marine environments, the remaining suborders preferred to reside in terrestrial ecosystems, particularly in soils, indicating a potential selectivity of geographical distribution. In conclusion, this study profiles a clear framework of diversity and distribution of the cosmopolitan myxobacteria and sheds light on the isolation of the uncultured myxobacteria.

**IMPORTANCE:** Myxobacteria are an attractive bacterial group ubiquitous in soil and aquatic environments. However, the biodiversity and ecological preferences of myxobacteria remain poorly understood across heterogeneous environments. We analyzed thousands of publicly available and high-quality 16S rRNA gene sequences of myxobacteria by using the phylogenetic reconstruction. The study presented an astonishing diversity than that expected from the previous studies. This study further demonstrated that the culturability of myxobacteria was perfectly comparable to its phylogeny in the phylogenetic tree. The geographical analysis clearly indicated that myxobacteria showed a nearly cosmopolitan distribution, while some taxa exhibited obvious preferences for specific environmental conditions. Together, our study provides novel insights into the diversity, distributions, and ecological preferences of of myxobacteria from diverse environments and lays the foundation for innovation of isolation techniques and the discovery of new secondary metabolites.

## INTRODUCTION

Myxobacteria are a large group of organisms, which taxonomically belong to the order *Myxococcales* within the class *Deltaproteobacteria* (1). They are one of nature’s “endowed” and extensively dispersed in natural environments, preferentially in places that are rich in microbes and organic matter. Myxobacteria are one of the fascinating and unique bacteria by virtue of their sophisticated social lifestyle and distinctive characteristics (2, 3), including cooperative swarming (4), group predation (5), multicellular fruiting body formation (6), and sporulation. Under favorable environmental conditions, vegetative cells of myxobacteria glide on the solid surface in swarm. According to the nutritional behavior and specialization in the degradation of biomacromolecules, myxobacteria are divided into two distinctive groups: micro-predators capable of lysing living microbial cells by excreting multiple lyases, and cellulose-decomposers (7). More notably, after the nutrition has been exhausted, most strains form species-specific and colorful fruiting bodies by oriented cell movement. Fruiting bodies contain a substantial number of desiccation-resistant myxospores, which can survive in hostile environments and are able to germinate under appropriate conditions even after decades of resting (8). A further intriguing feature of myxobacteria is the outstanding capability to produce structurally diverse bioactive secondary metabolites (9). Over the past decades, more than 100 new carbon skeleton metabolites and over 600 derivatives were identified from thousands of myxobacterial strains. These metabolites mainly comprise polyketides, non-ribosomal peptides, and hybrids thereof, and exhibit remarkable antifungal, antibacterial, cytotoxic, antiviral, immunosuppressive, antimalarial, and antioxidative activities with different modes of action (10). Therefore, myxobacteria have proven to be a promising source for new bioactive metabolites. In addition, some myxobacteria have been determined to lyse pathogenic bacteria and fungi (11), produce diverse carotenoids (12), degrade 2-chlorophenol (13), and reduce uranium(VI) (14), clearly demonstrating their wide applications in biomedicine, agriculture, industry, and environment.

As a phylogenetically coherent and distinctive group, members of the order *Myxococcales* have been currently divided into the three suborders mainly based on morphological properties of cells and sequences analysis of 16S rRNA gene, i.e., *Cystobacterineae*, *Sorangiineae*, and *Nannocystineae* (15). At the time of starting this study, the monophyletic order *Myxococcales* comprised 11 families, 29 genera, and 66 species enlisted in the List of Prokaryotic Names with Standing in Nomenclature database (LPSN, http://www.bacterio.net/, last accessed November 2018). In addition to a minority of type strains, majority of non-type strains have been isolated from the diverse ordinary and extreme habitats, and preserved in multiple culture collections or research teams, for example, more than 9000 myxobacteria collected by the group of Reichenbach and Höfle (16). Compared with the cultured myxobacteria, there are more the uncultured which were easily checked and identified via multiple culture-independent approaches such as clone libraries (17) and high throughput sequencing of 16S rRNA gene amplicons (18). These culture-independent approaches bypassed isolation and cultivation procedures and provided the information on in situ abundance and diversity of myxobacteria. Based on the numerous research findings, we inferred that myxobacteria in nature should have shown much higher diversity than what has been revealed so far. However, the true diversity of myxobacteria remains largely unknown until now, either because of the short amplicon sizes created or the not enough numbers of clone sequences analyzed in molecular surveys mentioned above.

Myxobacteria are widespread in terrestrial ecosystems (19). Historically and until recently, soils, decomposing plant materials such as tree bark, rotting wood, and leaves of trees, and dung of herbivores have always been considered as typical environments of these organisms (20). Sometimes, myxobacteria could have been isolated from limnetic habitats (18). According to the explanation for this phenomenon by Reichenbach (19), they are not actually indigenous limnetic organisms, and should originally come from terrestrial surroundings, and are finally transported into water bodies. Surprisingly, myxobacteria have also found in some extreme conditions, such as acidic wetlands (21), saline-alkaline soils (22), the arid desert (23), and even hot springs (24). More unexpectedly, a few myxobacteria have sporadically isolated from marine habitats (25), and different from known terrestrial myxobacteria, particularly in the requirement of sodium chloride roughly equivalent to the salinity of seawater (26). Thereafter, marine myxobacteria have attracted much attention worldwide. The study by Jiang *et al*. showed that myxobacteria-related sequences from four deep-sea sediments and a hydrothermal vent were diverse but phylogenetically similar at different locations and depths (17). They were separated from terrestrial myxobacteria at high levels of classification matching a phylogenetic distance of approximately 10%, likely resulting from geographic separation and environment selection (17). Soon after, Brinkhoff *et al*. demonstrated that an obligate marine myxobacteria cluster (MMC) constituted up to 13% of total bacterial 16S rRNA genes in surface sediments of the North Sea, and was also detected in most sediment samples from other sea areas (27). These brief statements described above suggest that unique habitats may harbor specialized taxa of myxobacteria. We hypothesized that myxobacteria could be more taxonomically diverse than what is acknowledged presently, due to the enormously diverse distinct niches at global scale. However, the detailed relationships between more taxa and different environments and distribution patterns of myxobacteria have not yet been thoroughly investigated.

With the development of next-generation sequencing technologies and the increasing studies on myxobacteria, thousands of 16S rRNA gene sequences of myxobacteria have been rapidly accumulated in public databases. These available sequences thus provide an ideal opportunity for attempting to answer the above-mentioned questions. In the current study, we first collected and integrated 16S rRNA gene sequences from the multiple databases. The phylogenetic analysis based on myxobacteria-related sequences was then performed to infer their taxonomic diversity. Using this extensive survey, we addressed the following questions: (i) how many taxa of the cultured and uncultured myxobacteria at different taxonomic levels are there so far? (ii) what is the environmental distribution pattern of myxobacteria inhabiting diverse environments worldwide? and (iii) what are the ecological preferences of individual myxobacterial lineages?

## RESULTS

### The outline of 16S rRNA gene sequences of myxobacteria

The 16S rRNA gene sequences used in this study were from PCR amplicons, clones, genomes, and metagenome assemblies, respectively. For isolation sources, sequences from the soil, marine organism, marine sediments, activated sludge, and terrestrial organism accounted for 59.42%, 9.11%, 6.92%, 6.80%, and 5.08%, respectively, while the remaining sequences from others only contributed 12.67% (Table S1 and Table S3). The isolation locations of some sequences with the exact longitudes and latitudes were marked by red dots with black rings in Figure 1, most of which were situated in East Asia, South Asia, Europe, and North America. The sequences were obtained from more than 80 countries and areas (Figure 1 and Table S4). More than 80% of sequences were from ten countries, including China, Denmark, the USA, Mexico, Panama, Japan, India, Germany, Spain, and France. A small number of sequences came from the oceans, such as the Pacific Ocean and the Atlantic Ocean. Therefore, all the sequences in this study were not only in huge numbers, but also from extensive sources including diversified habitats and many countries/areas, adequately demonstrating the broad representativeness of sequences.

**Figure 1.**
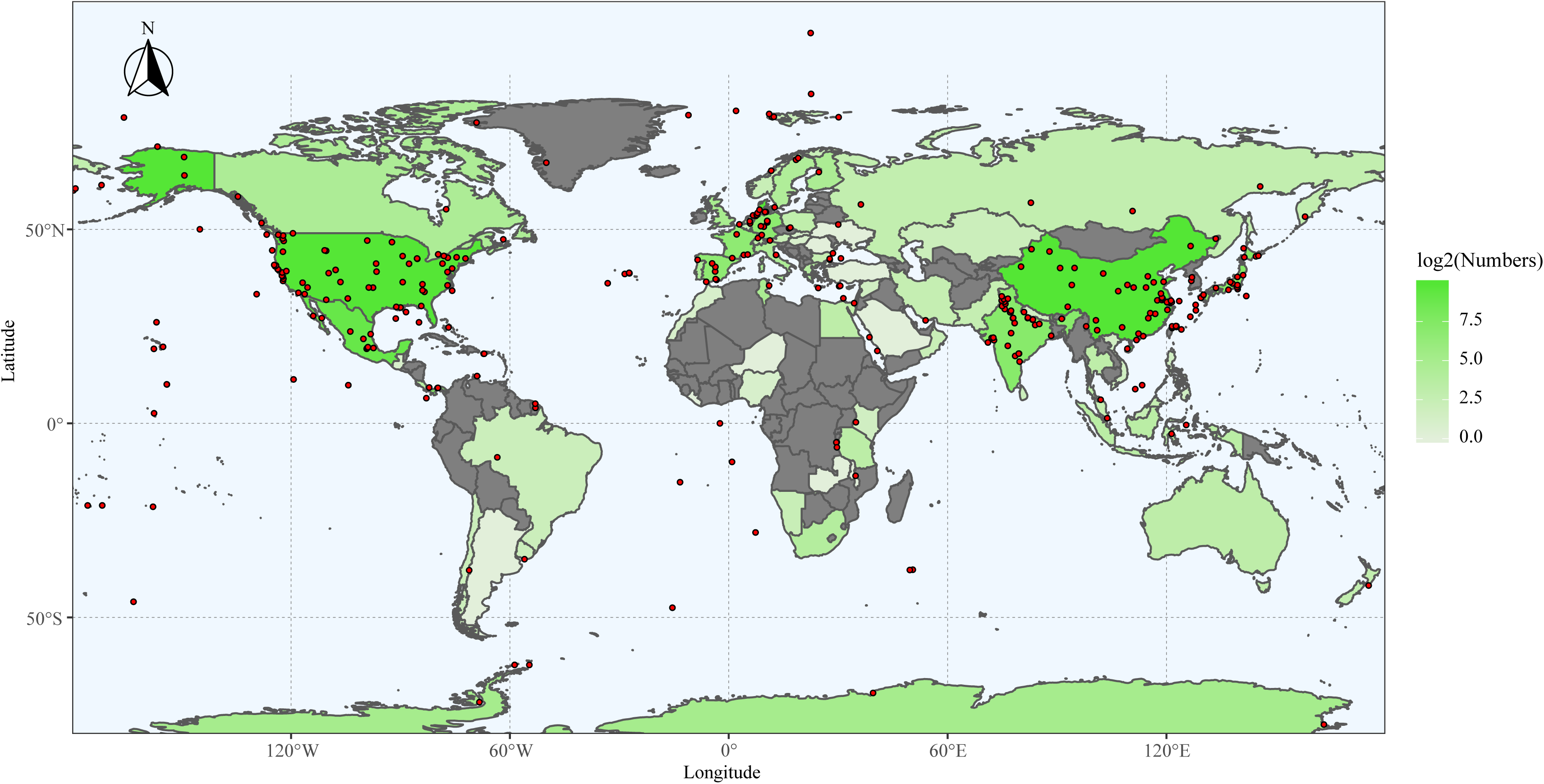
Map of the geographical distribution of the strains showing the isolation locations marked by red dots with black rings, the countries, and areas. The countries and areas were marked by gradient colors from “#E2EFDA” to “#54E635” based on the logarithmic values of sequence numbers. Some countries and areas with a few sequences were not illustrated in the map and were shown in Table S1. The graph was plotted by using the “ggplot2” package with the geom.sf() function (https://cran.r-project.org/web/packages/ggplot2/index.html) according to the longitude and latitude coordinates of isolated sites. The world map was generated by using the “rworldmap” package (https://cran.r-project.org/web/packages/rworldmap/index.html).

### The unexpected diversity of myxobacteria

A highly resolved phylogenetic tree of 16S rRNA gene sequences clearly presented an unexpected taxonomic diversity of myxobacteria (Figure 2 and Fig. S1). Specifically, myxobacteria were divided into 20 suborders, 58 families, 445 genera, and 998 species according to phylogenetic positions and taxonomic criteria mentioned above (Table 1). At the suborder level, 20 suborders were comprised of the three previously described suborders (*Cystobacterineae*, Sorangiineae, and *Nannocystaceae*) (28) and 17 newly discovered suborders (Suborder_1 to Suborder_15). In this study, inspired by definitions of abundant and rare species (29), the abundant and rare suborders were defined based on the proportion of 16S rRNA gene sequences, with ≥ 0.5% as abundant suborder and < 0.5% as rare suborder. Under the criterion, abundant suborders included *Sorangiineae*, *Cystobacterineae*, *Nannocystaceae*, Suborder_16, Suborder_15, Suborder_17, and Suborder_4, accounting for 29.08%, 26.58%, 20.89%, 10.67%, 4.72%, 4.14%, and 2.88% of all sequences, respectively (Table 1). Rare suborders contained the remaining thirteen taxa, namely, Suborder_1 to Suborder_3 and Suborder_5 to Suborder_14, accounting for 1.04% of all sequences altogether. Obviously, abundant suborders showed significant overrepresentation relative to the thirteen rare suborders.

**Figure 2.**
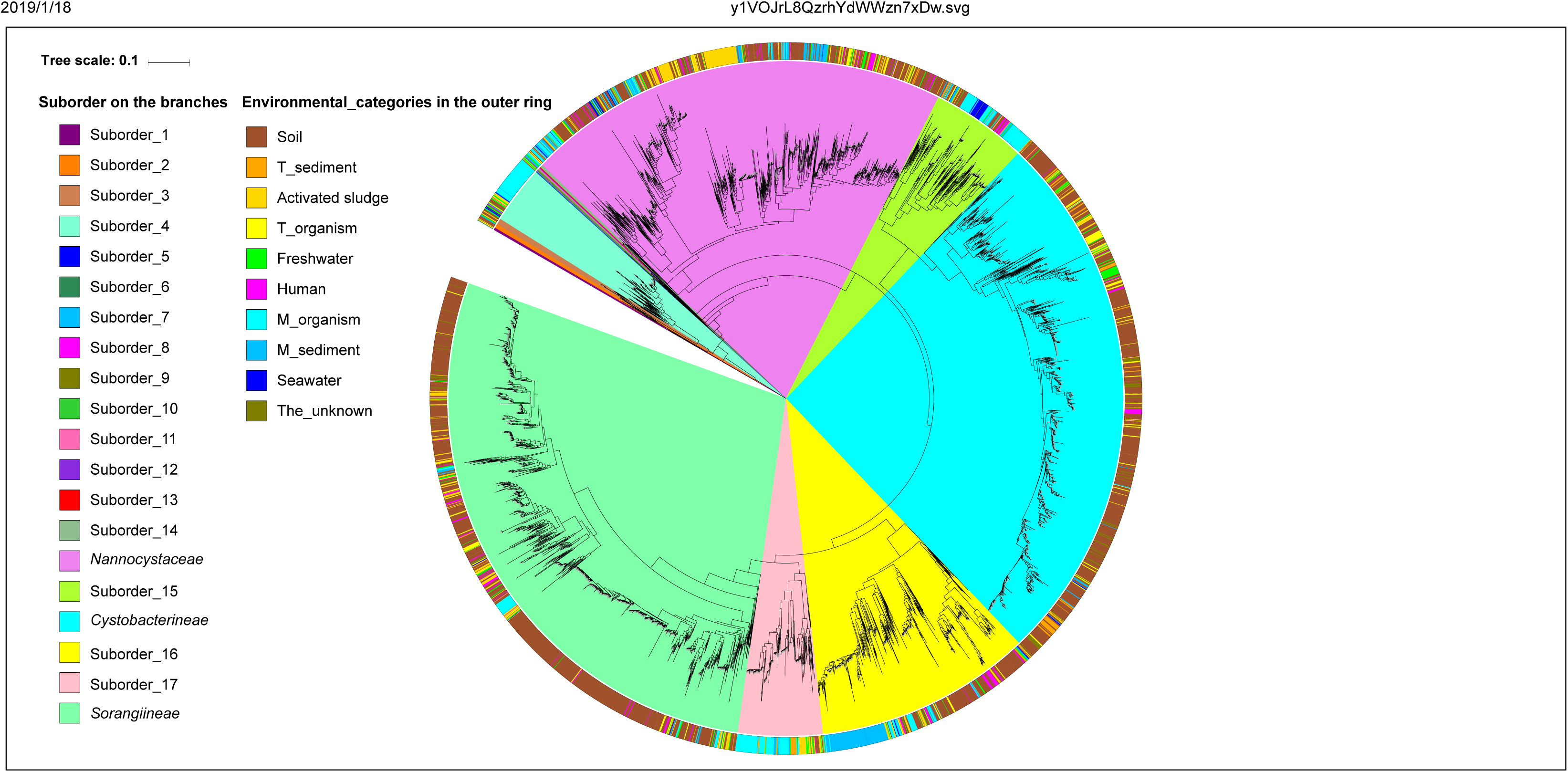
The phylogenetic tree inferred from 16S rRNA gene sequences showing the positions of bacteria within the order *Myxococcales*. The sequence of *Desulfovibrio desulfuricans* ATCC 27774 roots the tree. Scale bars: 0.1 substitutions per nucleotide position. Suborder-level clusters are indicated by 20 different colors (as shown in Table S2) on the branches of the tree, and environmental categories of all sequences are indicated by ten different colors in the outer ring of the tree.

**Table 1.** The detailed taxonomy of myxobacteria at the four taxonomic levels

| Abundant<br>/ Rare | Suborder | Family | Genus | Species | Numbers<br>(proportion, %) |
| --- | --- | --- | --- | --- | --- |
| Rare | Suborder_1 | 1 | 5 | 5 | 5 (0.10) |
| Rare | Suborder_2 | 1 | 3 | 6 | 10 (0.20) |
| Rare | Suborder_3 | 1 | 3 | 8 | 15 (0.30) |
| Abundant | Suborder_4 | 1 | 28 | 44 | 144 (2.88) |
| Rare | Suborder_5 | 1 | 1 | 1 | 1 (0.02) |
| Rare | Suborder_6 | 1 | 1 | 1 | 1 (0.02) |
| Rare | Suborder_7 | 1 | 1 | 1 | 1 (0.02) |
| Rare | Suborder_8 | 1 | 1 | 1 | 2 (0.04) |
| Rare | Suborder_9 | 1 | 2 | 2 | 2 (0.04) |
| Rare | Suborder_10 | 1 | 1 | 1 | 1 (0.02) |
| Rare | Suborder_11 | 2 | 3 | 3 | 4 (0.08) |
| Rare | Suborder_12 | 1 | 1 | 1 | 1 (0.02) |
| Rare | Suborder_13 | 1 | 2 | 2 | 2 (0.04) |
| Rare | Suborder_14 | 2 | 3 | 3 | 7 (0.14) |
| Abundant | *Nannocystaceae* | 11 | 157 | 309 | 1044 (20.89) |
| Abundant | Suborder_15 | 10 | 48 | 108 | 236 (4.72) |
| Abundant | *Cystobacterineae* | 6 | 50 | 131 | 1328 (26.58) |
| Abundant | Suborder_16 | 4 | 54 | 116 | 533 (10.67) |
| Abundant | Suborder_17 | 6 | 32 | 60 | 207 (4.14) |
| Abundant | *Sorangiineae* | 5 | 49 | 195 | 1453 (29.08) |
| Total | 20 | 58 | 445 | 998 | 4997 |

At the family level, the dominant families constituting above 10% of all sequences were Cystobacterineae_Family_6 (21.83%), Sorangiineae_Family_5 (16.75%), and Sorangiineae_family_4 (12.13%); the subdominant families constituting above 5% were Nannocystaceae_Family_11 (5.74%) and Suborder_16_Family_4 (5.66%); the remaining 53 families contained relatively few myxobacteria-related sequences (Table S5). At the genus level, the Cystobacterineae_Family_6_Genus_27 (15.47%) was the most dominant genus; the Sorangiineae_Family_4_Genus_21 (6.80%) and Sorangiineae_Family_5_Genus_19 (4.62%) were the most second and third dominant genera; each remaining genus constituted less than 3% of total sequences (Table S5). At the species level, Cystobacterineae_Family_6_Genus_27_Species_5 (12.55%), Sorangiineae_Family_4_Genus_21_Species_1 (6.80%), and Sorangiineae_Family_5_Genus_19_Species_3 (4.54%) were dominant species relative to other non-dominant species (each less than below 3%, Table S5). These results indicated that myxobacteria presented an incredible diversity than that expected from the previous studies and were also greatly overrepresented by several dominant taxa at taxonomic levels from family to species.

The comparison between the numbers of taxa identified in this study and those previously identified at each taxonomic level were conducted. According to the newly rigorous criteria above, the validly described type strains of the order *Myxococcales* were reclassified into four suborder, seven families, 13 genera, and 24 species (Table S6). Therefore, we concluded that 16 new suborders, 51 new families, 432 new genera, and 974 new species were determined in this study. Meantime, the occurrence frequencies (%) of the new family, genus, and species within the three known suborders were compared based on the sufficiently clear taxonomic determination of myxobacteria. As shown in Table S7, abundant new taxa at taxonomic levels from family to species were found among the three suborders. High occurrence frequencies of new taxa were impressive, especially those of new genus and species above 90%. The rarefaction is one of the most widely used methods to assess species richness from the results of sampling. Here, we constructed two types of rarefaction curves for all 16S rRNA gene sequences based on abundant suborders and environments. The sharp differences of sequence numbers among the “samples” are not suitable for data normalization and further comparative analysis. Therefore, we only focused on the trend of each rarefaction curve. Both suborder-based and environment-based rarefaction curves were far from the plateau at 97% identity (Figure 3), clearly indicating that more species could be found in either each suborder or each environmental category as more sequences are checked. The patterns were also supported by the coverage values of “samples” (Table S8). Therefore, the results explicitly demonstrated that the currently defined taxa of the order *Myxococcales* are just the tip of the iceberg.

**Figure 3.**
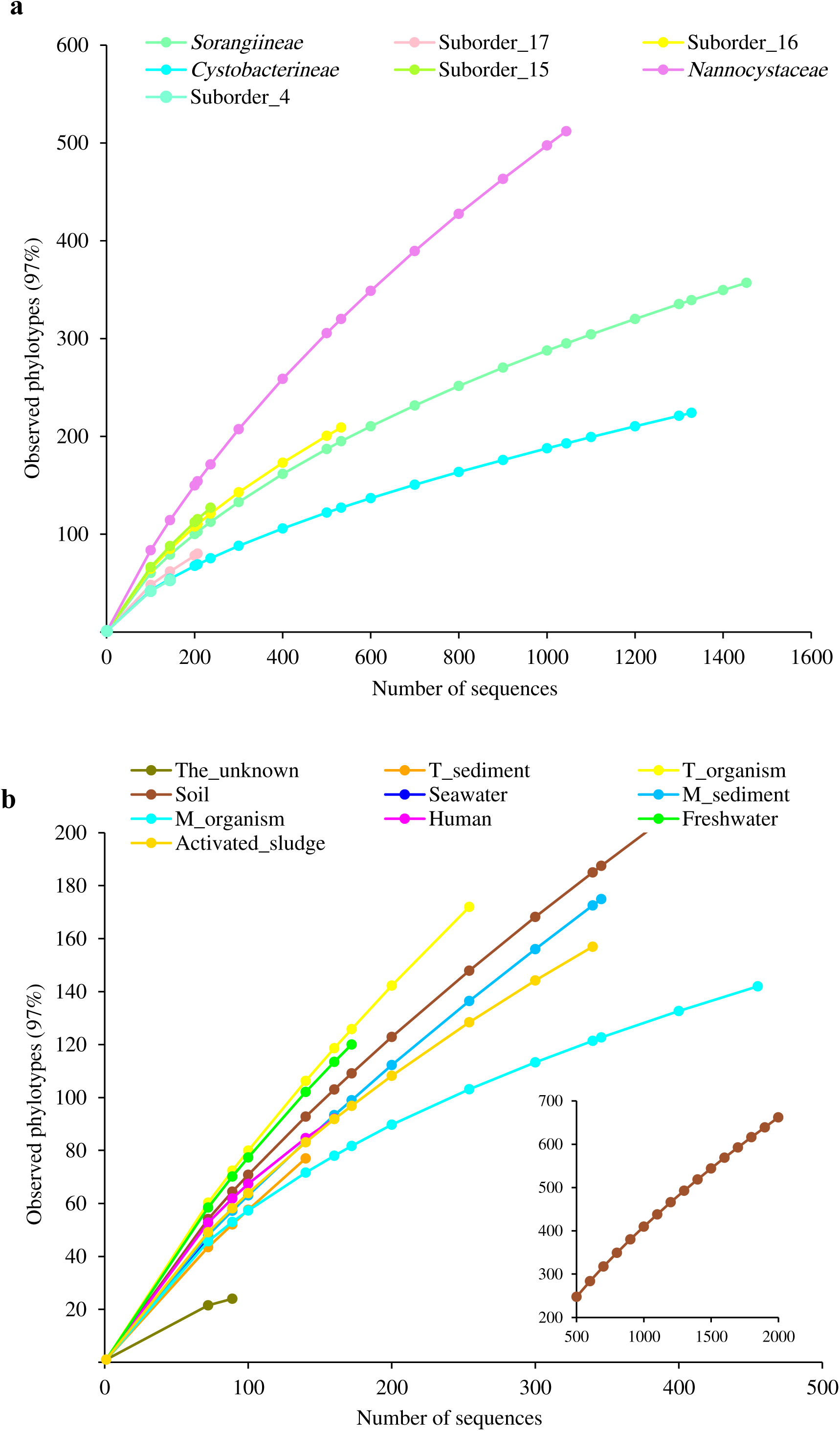
Rarefaction curves for all 16S rRNA gene sequences based on the seven abundant suborders (a) and ten environmental categories (b). Operational taxonomic unit s was determined at 97% identity. The colors of points and curves in the figure were listed in Table S2.

### The expanded phylogeny of myxobacteria

After the determination of the taxonomic diversity of myxobacteria, the evolutionary relationships were also inferred from the phylogenetic tree of 16S rRNA gene sequences. As shown in Fig. S2, myxobacteria consisted of three well-supported lineages, designated as lineage A, lineage B, and lineage C. More specifically, the minor lineage A congruent with the Suborder_1 was phylogenetically distant from the other two lineages, and located at the base of the tree, theoretically suggesting that lineage A should be a common ancestor of all descendants of myxobacteria. However, it is surprising that the five strains of lineage A resided in distinctive habitats located in different geographic regions, namely deep-sea sediment of the Pacific Ocean, soil in Brazil, periphyton in China, rice paddy field soil in Japan, and cave water in the USA. The lineages B and C were more closely related to each other than the lineage A and thus were considered as sister lineages. The medium lineage B holding 169 myxobacteria-related sequences was on the equivalent of the three suborders from Suborder_2 to Suborder_4. The largest lineage C consisted of six abundant suborders and ten rare ones. In lineage C, the Suborder_5 occupied the basal-most position; the next five suborders from Suborder_6 to Suborder_14, which totally contained 22 strains, were later and sub-basal taxa; the remaining six suborders were phylogenetically young taxa based on their position and dominant taxa according to the high proportion of 96.1% of all sequences.

Within these isolates, there were 3890 uncultured myxobacteria, 1101 cultured ones, and 6 ones with few indications whether they have been cultured or not. With an exception of type strain DSM 53668^T^ (CP011125), all cultured strains were affiliated to the three known suborders, which was indicative of their easy-to-culture characteristics, and vice versa for uncultured strains. For isolation environments from soil to human, as shown in Fig. S3, the percentages of the cultured decreased gradually, and conversely, those of the uncultured increased successively, implying that the expected probabilities of finding the new genotypes or strains from the latter environments are much greater than those from the former. More intriguingly, we observed that cultured myxobacteria including the well-described type strains were overall inclined to locate in the shallow branches of the phylogenetic tree; on the contrary, those of the uncultured myxobacteria the deep branches (Fig. S1). Therefore, the culturability of myxobacteria was perfectly comparable to the distribution patterns of branches in the 16S rRNA-based phylogenetic tree.

### Global biogeographic distribution of myxobacteria

To determine the geographical distribution patterns of myxobacteria, the suborder-level sequences were summarized and compared according to isolation environments. As shown in Figure 1 and Table S1, myxobacteria were widely distributed in both terrestrial and marine environments, thereby suggesting a cosmopolitan distribution in these habitats. Considering a small number of sequences, the distribution of the rare suborders was no longer analyzed in depth. The myxobacteria within each abundant suborder contained all environmental categories. Nevertheless, as shown in Figure 4a, they were subjected to uneven distributions across environments to some degree. More specifically, *Sorangiineae*, *Cystobacterineae*, and *Nannocystaceae* were mainly from terrestrial environments, especially from the soil. On the contrary, Suborder_4, Suborder_15, and Suborder_17 were dominant in marine environments, including marine organism, marine sediments, and seawater. Moreover, the distribution of sequences at the family level was also unequal in multiple environments, as illustrated in Figure 4b. The results shed light on possibility to isolate myxobacteria of specific taxa from specific habitats. For example, we are more likely to isolate strains of Family_5 and Family_6 in Suborder_17 from the hypersaline mat (Table S1), which was belonged to the M_organism. Strikingly, the two previously neglected environments including activated sludge and human skin also inhabited a certain number of myxobacteria within each abundant suborder. Furthermore, more than half of myxobacteria from activated sludge belonged to the suborder *Nannocystaceae*. Contrariwise, myxobacteria from human skin scattered over the seven abundant suborders.

**Figure 4.**
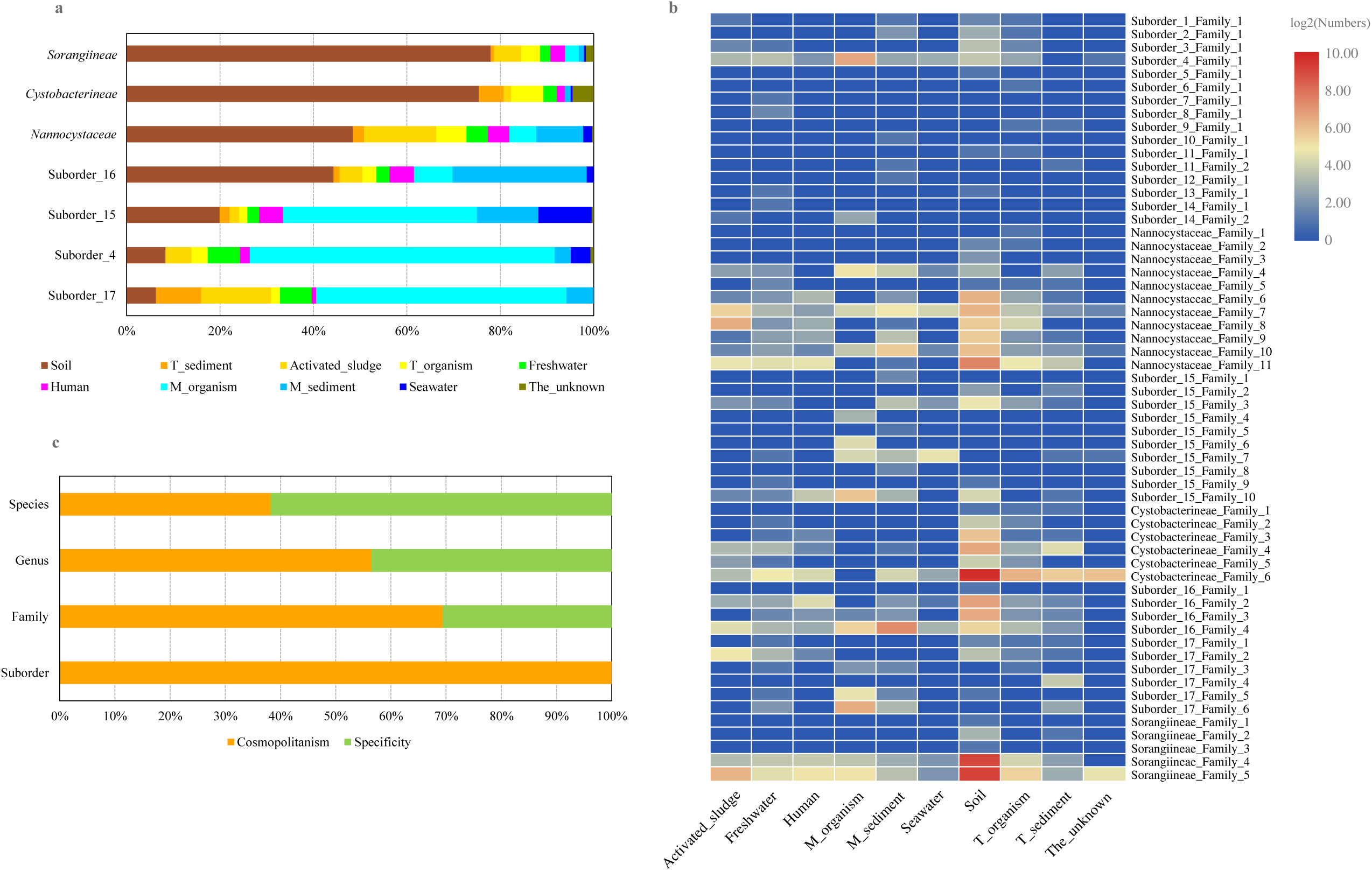
The distribution of the seven abundant suborders (a) and 58 families (b), and the proportion of specific and cosmopolitan taxa at the four taxonomic levels (c).

A comprehensive analysis of the relationships between myxobacterial taxa mainly including generalists (ubiquitous, cosmopolitan taxa) and specialists (environment-specific taxa) and different environments at the four taxonomic levels was performed. In this study, we defined a specificity criterion as having 80% or more of their observations (≥5) of single taxon belonged to a single environmental category, otherwise, it was defined as a cosmopolitanism. As illustrated in Figure 4c, the specificity of myxobacteria showed an increasing trend from the suborder to species levels, whereas cosmopolitanism a reverse trend. It is also worth noting that the specificity of taxa is closely related to environmental categories and taxa number. With the number of the environmental categories or taxa increasing, specificity may decrease or disappear, even at species level. Certainly, according to the current criterion, habitat specificity of some taxa at family, genus and especially species levels were evidently shown in this study (Figure 1, Table S1, and Table S9), including the known MMC cluster matching the genus Suborder_16_Family_4_Genus_33 from marine sediments.

## DISCUSSION

Compared to many in-depth studies on the physiological basis of social behavior and on action mechanisms of secondary metabolites of myxobacteria over the last several decades, much less attention has been hitherto paid to their taxonomic diversity and biogeographic distribution across heterogeneous environments. In this case, by using the public massive 16S rRNA gene sequences of myxobacteria, we performed a comprehensive and systemic research to expand our understanding of the two aspects above. These findings demonstrate that myxobacteria from various environments have remarkable taxonomic diversity and widespread geographic distribution, which is far beyond what is expected from previous studies.

From the discovery of myxobacteria to the present, an increasing number of bacteria has been isolated by using different media and culturing conditions. In the course of the study on the correlations between morphological characteristics and 16S rRNA gene phylogeny of myxobacteria, the three lineages/suborders (*Cystobacterineae*, *Nannocystineae*, and *Sorangiineae*) were determined via branching pattern of the phylogenetic tree (15). About ten years later, the study by Garcia *et al*. not only reaffirmed the three suborders of myxobacteria, but also discovered nine new taxa, which were probably at family level, based on the expanded phylogeny of 16S rRNA gene sequences of 101 myxobacteria (28). With the rapid development of sequencing technology, the growing number of uncultured myxobacteria have been detected by using culture-independent methods, such as the high-throughput 16S rRNA amplicon sequencing and metagenomic sequencing. Among these ecological surveys, the great diversity of myxobacteria were documented in the varied ecological niches (17, 18), and more meaningfully, some new habitat-specific clusters were found, such as the MMC cluster as mentioned above (27, 30, 31). However, from the perspective of classification, the real phylogenetic diversity of cultured and uncultured myxobacteria is still unknown, and the accurate taxonomic affiliation of most new taxa has not yet been determined to date. Here, the present study demonstrates that thousands of myxobacteria are highly diverse at the four taxonomic ranks from species to suborder, most members of which are defined as novel taxa based on the 16S rRNA gene phylogenetic reconstruction. Moreover, we provided a relatively complete and stable evolutionary framework of myxobacteria for classification of currently described new taxa and subsequent cultured and uncultured strains, especially those distantly related to valid type strains. Just recently, the positive correlation between taxonomic distance and the production of distinct secondary metabolite families was substantiated through the systematic metabolite survey of about 2300 myxobacteria (16), which provides an important strategic implication that the use of bacteria belonging to new species, genera, families, and even suborders could substantially increase the discovery chance of new natural products, and further highlights the importance of the myxobacterial classification system within this scenario. Therefore, a more detailed taxonomic system of myxobacteria established in the current study could fuel the discovery of interesting novel compounds for the foreseeable future.

Similar to actinobacteria and bacillus, myxobacteria have been considered for a long time as typical terrestrial organisms (20). By now, along with the more phylogenetic analyses of myxobacteria from marine environments, it has been widely accepted that some distinctive marine myxobacteria are widespread in upper oxic sediments and bottom anoxic sediments from different locations and depths (17, 27), which are phylogenetically distinct from soil and limnetic bacteria (32). In addition to natural marine sediments (33), myxobacteria as the second abundant group also presented in oil-polluted subtidal sediments, with their relative abundance decreasing through depth (34). Somewhat incomprehensibly, a few myxobacteria are only sporadically found in oxic seawater samples (27), which is consistent with our result of the presence of only about 1% seawater bacteria. On the contrary, Ganesh *et al*. described the metabolically active myxobacteria representing up to 3% of total sequences in the larger size fraction from a marine oxygen minimum zone (35). These results indicated that strict or facultative anaerobic myxobacteria being little concerned for a long time are also dominant in marine environments, such as the well-described aerobic relatives. More studies about how myxobacteria live in oxygen-free environments and what the metabolic differences between aerobes and anaerobes are needed to be further elucidated.

Meanwhile, some myxobacteria reside in less focused environments, such as activated sludge (36, 37) and microbial mats/biofilm (38). In the study by Wang *et al*. (39), the order *Myxococcales* is one of the core orders of microbial community with a range from 0.86 to 9.5% of the classified sequences from activated sludge samples. But, the role of myxobacteria in these biological wastewater treatment systems has not been well studied. Besides, unexpectedly, more than 3% of myxobacteria in this study were from the skin of children with atopic dermatitis (40, 41). Therefore, we call upon all myxobacteriologists that more attention should be paid to the impact of myxobacteria on human and animal health in subsequent studies, although no pathogenic myxobacteria have been observed up to now. Altogether, consistent with the report by Tamames *et al*. (42), the two possible geographical patterns for myxobacteria have been determined in this study: most myxobacteria at high taxonomic ranks of family and suborder could share many environments; some at the species and genus levels could be found specifically within each environment. The underlying adaptation mechanisms for pandemic and endemic distribution patterns of myxobacteria under different taxonomic hierarchy remain to be explored.

At present, the isolation, purification, and cultivation of myxobacteria are one of the biggest challenges, which are still essential for elucidating myxobacterial physiology in depth. Numerous strategies have been successfully used to increase the cultivation efficiency of myxobacteria as well as to isolate novel taxa, for example, isolation of salt-tolerant myxobacteria from marine conditions (25). There is no doubt that the present study may provide preliminary clues for the culture of previously uncultured members of myxobacteria. On the one hand, after determining the biogeographic distribution of myxobacteria, we can select environmental samples purposefully rather than randomly and blindly to isolate target bacteria, despite that isolation efforts may be in vain. For example, when the MMC is targeted, the chance of successful isolation from marine sediments is much greater than that from other habitats (27). Based on this study and those by others, the previously neglected habitats with an astonishing diversity of myxobacteria are promising reservoirs for getting yet-to-be-cultured bacteria. On the other hand, our study clearly indicates that isolation efforts under anaerobic conditions may be a more effective alternative for obtaining myxobacteria (43). The characterization and description of the species *Anaeromyxobacter dehalogenans* as the first facultative anaerobic myxobacteria have proven the feasibility of anaerobic cultivation approaches (13). Additionally, based on the fact of the MMC restricted to salinities ranging from 6 to 60 practical salinity units (27), salinity is another very important factor and thus should be paid more attention in the process of isolating myxobacteria.

In this report, we contributed a far more comprehensive knowledge to the expanded diversity and biogeographic distribution pattern of myxobacteria from diverse environments, which was based on the phylogenetic reconstruction of thousands of 16S rRNA gene sequences. The myxobacteria analyzed in this study are considerable diverse with a wide distribution in terrestrial and marine environments, and harbor abundant new taxa at each taxonomic rank. This study structures a robust taxonomic framework of myxobacteria and provides overarching guidance for innovation of isolation techniques and the discovery of new secondary metabolites.

## MATERIALS AND METHODS

### Dataset construction of 16S rRNA gene sequences of myxobacteria

A considerable number of 16S rRNA gene sequences of myxobacteria with various lengths have been accumulated in multiple public databases including NCBI database, SILVA database (44), RDP database (45), and Integrated Microbial Genomes (IMG) database (46). We retrieved 16S rRNA gene sequences from the four databases above. For NCBI database, a preliminary search using the keyword of “Myxococcales [ORGANISM] AND 16S [TITLE]” in the nucleotide database was performed. To obtain enough genetic information and to embrace more available sequences, near-full-length (≥1200 bp) sequences (hereafter defined as Dataset 1, the same below) were picked out from candidate items. Using 16S rRNA gene of type strain *Myxococcus xanthus* DSM 16526^T^ (accession number: DQ768116.1) as a query sequence, sequences from the complete and draft genomes of bacteria within the order *Myxococcales* (Dataset 2) were obtained by using blast genome alignment (90% coverage and 78% identity) with default algorithm parameters. Both multiple-copies and single-copy sequences from the respective genome were used in the following analysis. Thousands of sequences (Dataset 3) were searched and downloaded according to their taxonomic context in SILVA database (release 132). The target sequences (Dataset 4) were obtained from RDP database under these options including the type and non-type strains, uncultured and cultured isolates, with the sequence length at or above 1200 bp, good quality, and nomenclatural sequences. The sequences (Dataset 5) were acquired from IMG database by using the BLAST databases including 16S rRNA public isolates and public assembled metagenomes with an E-value of 1e-5 on Oct. 29th, 2018. All sequences from the five databases were integrated as a raw dataset.

### Sequences validation and environmental categories of myxobacteria

The sequence validation was carried out according to the following rigorous criteria: (1) the longest high-quality sequence was selected as the representative of the same strain with different original numbers; (2) the ambiguous base ratio of each sequence was less than 0.2%; (3) the sequences which do not belong to the order *Myxococcales* were excluded based on the screening of RDP Classifier with more than 80% confidence; (4) chimeric sequences were identified by the free package USEARCH61 (47) and then removed. Finally, a total of 4997 high-quality 16S rRNA gene sequences including those of all type strains were analyzed with the GenBank accession numbers and IMG gene IDs indicated in Table S1. The definition of the cultured and uncultured myxobacteria was according to user-provided metadata or literature retrieval. Meantime, the isolation sources of all sequences were retrieved from the original annotations, published studies, or documents of culture collections, and then manually assigned to multiple defined environmental categories referring to the previously described schema (42) (48). To simplify categorizations, sequences from marine animals, marine plants, marine biofilms, and others were classified as sequences from marine organisms, whereas those from terrestrial animals, terrestrial plants, terrestrial biofilms, and others as sequences from terrestrial organisms. Notably, sequences from activated sludge and human skin, although in terrestrial ecosystems, were designedly divided into two different categories, in order to highlight their uniqueness and distinctiveness. Altogether, ten environmental categories were designated as Marine_organism (hereafter abbreviated as M_organism, the same below), Marine_sediment (M_sediment), Seawater, Soil, Terrestrial_sediment (T_sediment), Activated_sludge, Terrestrial_organism (T_organism), Freshwater, Human, and The_unknown (the unknown isolation sources), as shown in Table S1. The isolation geographical coordinates and countries of sequences were also determined according to user-provided metadata or literature retrieval as much as possible. Rarefaction curves of all samples were calculated by using the software Mothur version 1.38 (49).

### Phylogenetic analysis of 16S rRNA gene sequences of myxobacteria

To analyze the phylogeny of 16S rRNA gene, all sequences were aligned by using the software MAFFT version 7.037b with default settings (https://mafft.cbrc.jp/alignment/software/) (50). The approximately maximum-likelihood (ML) phylogenetic tree of 16S rRNA gene sequences was drawn by using the JTT+CAT model and 1000 bootstrap replicates within the software FastTree version 2.1.10 (51). In order to make the display more intuitive and concise, bootstrap values were only indicated by solid circles filled by light blue (rgba(200, 200, 255, 0.8)) with different sizes ranged from 0.5 px to 3 px on the middle (50%) of branches of the phylogenetic tree in supplementary Fig. S1. To compare with previous studies, the sequence of *Desulfovibrio desulfuricans* ATCC 27774 (accession number: M34113), a sulfate-reducing bacterium also in the class *Deltaproteobacteria* (52), was chosen as an outgroup to root the phylogenetic tree. The visualization, annotation, and management of phylogenetic tree were performed by using the web-based tool of Interactive Tree Of Life (iTOL) (53). Meantime, the taxonomic hierarchy of all sequences was determined based on the strict criteria below: the identity of 97.0% or lower for two 16S rRNA gene sequences serves as strong evidence for distinct species, 94.5% or lower for distinct genera, 89.0% or lower for distinct families, and 85.0% or lower for distinct suborders (28, 54, 55). For the sake of comparison, the taxonomic affiliation of all type strains was re-identified based on the above criteria. The pairwise identities of aligned sequences were calculated by using the software DNAMAN version 8 (Lynnon Biosoft, USA, https://www.lynnon.com/) with the distance method of Kimura. Combined with phylogenetic analysis and the above taxonomic thresholds, the classification of each sequence was carried out with the artificial modification, if necessary. In addition, in order to ensure the repeatability and reproducibility of all figures, the Hex values and rgb/rgba values of colors used to mark different elements including suborders, environmental categories, the culturability, lineages, etc. were also provided, as shown in Table S2.

### Data accessibility

All 16S rRNA gene sequences were retrieved from public databases. Additional data are all provided as Supplementary datasets in Supplementary materials. For more details, the availability is held by the corresponding authors upon request.

## SUPPLEMENTAL MATERIAL

Supplemental material for this article may be found at https://doi.org/XXXX.

## ACKNOWLEDGMENTS

We gratefully acknowledge Dr. Linfeng Gong at Third Institute of Oceanography (Xiamen, China) for the generous help in analysis for rarefaction curves.

Funding for this work was provided by the GDAS’ Special Projects of Science and Technology Development [2019GDASYL-0401002], the National Natural Science Fund of China (31800003 and 31600008), the Guangdong Province Science and Technology Innovation Strategy Special Fund (2018B020205003), and the Science and Technology Plan Project of Guangdong Province [2019B030316010].

